# Visualizing subcellular structures in neuronal tissue with expansion microscopy

**DOI:** 10.1101/2020.12.05.408724

**Authors:** Logan A. Campbell, Katy E. Pannoni, Niesha A. Savory, Dinesh Lal, Shannon Farris

**Author notes:** Authors contributed equally (Logan A. Campbell, Katy E. Pannoni).

## Abstract

Protein expansion microscopy (proExM) is a powerful technique that crosslinks proteins to a swellable hydrogel to physically expand and optically clear biological samples. The resulting increased resolution (~70 nm) and physical separation of labeled proteins make it an attractive tool for studying the localization of subcellular organelles in densely packed tissues, such as the brain. However, the digestion and expansion process greatly reduces fluorescence signals making it necessary to optimize ExM conditions per sample for specific end goals. Here we describe a proExM workflow optimized for resolving subcellular organelles (mitochondria and the Golgi apparatus) and reporter-labeled spines in fixed mouse brain tissue. By directly comparing proExM staining and digestion protocols, we found that immunostaining before proExM and using a proteinase K based digestion for 8 hours consistently resulted in the best fluorescence signal to resolve subcellular organelles while maintaining sufficient reporter labeling to visualize spines and trace individual neurons. With these methods, we more accurately quantified mitochondria size and number and better visualized Golgi ultrastructure in reconstructed CA2 neurons of the hippocampus.

## INTRODUCTION

Protein retention expansion microscopy (proExM) is a powerful tool that crosslinks proteins to a swellable hydrogel to optically clear and physically expand tissues up to ~4-fold in volume^1,2^. Because expansion physically separates crosslinked moieties, this technology is particularly useful for visualizing subcellular structures in densely packed tissues, such as the brain. However, one consequence of tissue expansion is a decrease in the fluorescence intensity of labeled proteins primarily due to the digestion process and the dilution of fluorescence signal per unit volume. Various ExM protocols have described different methods for improving fluorescence retention, primarily by modifying fixation, crosslinking, and/or digestion conditions to preserve protein epitopes^2–8^. One common ExM protocol uses a strong protease-based digestion (proteinase K^2^), but other gentler proteases have also been used (LysC^2,3^), as well as a combination of heat and detergents in place of proteases (e.g. autoclave in an alkaline buffer^2–5^).

Immunostaining can also be done before or after ExM to boost fluorescence^2,3^ (Fig. 1), although results are often dependent on the protein epitope and the quality of antibody staining. To improve antibody staining in brain tissue, antigen retrieval is often performed prior to immunostaining via boiling tissue in water or heating tissue in a citrate buffer (pH 6) to expose protein epitopes and reduce nonspecific staining. Here we set out to compare ExM immunostaining and digestion conditions for fluorescently labeled subcellular organelles in perfused brain sections using antibodies that either require or do not require antigen retrieval (COX4-labeling of mitochondria and GOLGA5-labeling of Golgi, respectively). Visualizing the spatial organization of organelles within compartmentalized cells, such as neurons, is important for understanding their function, thus we paid particular attention to conditions that maintained fluorescence signal from genetically-encoded reporters (i.e. enhanced green fluorescent protein, EGFP, and tdTomato). We found that performing IHC, with or without antigen retrieval, prior to ExM with proteinase K digestion for 8 hours best preserves fluorescence signal for co-visualizing subcellular organelles and neuronal morphology, including spines. Further, we report our optimized conditions for antibodies against widely used reporters and conclude with protocol considerations for achieving specific end goals.

**Figure 1:**
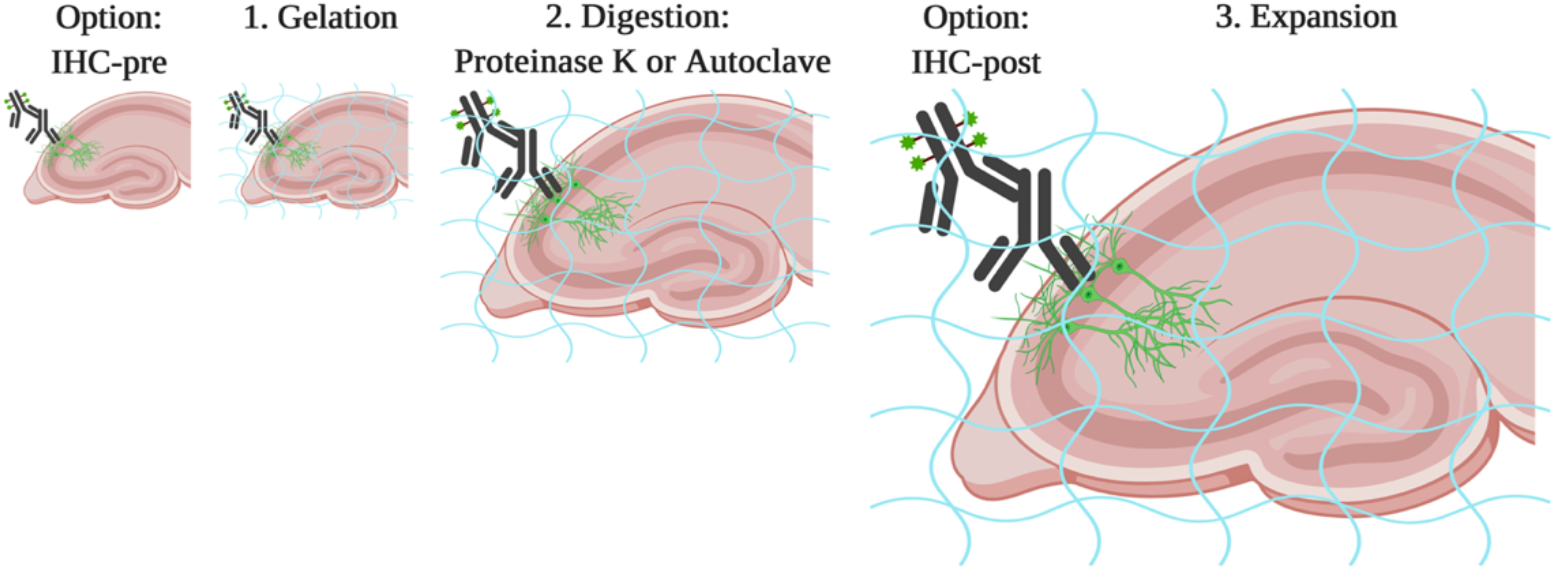
ExM workflow for visualizing subcellular organelles in fixed brain tissue. Schematic of immunostaining and proExM workflows tested to optimize visualization of organelles in genetically-labeled neurons.

## RESULTS

### Proteinase K digestion time impacts fluorescence intensity and expansion factor

Sufficient digestion or homogenization (also referred to as hydrolysis in some protocols) is required to prevent sample distortion during expansion and is highly dependent on digestion conditions, including time, temperature, pH, buffer composition and enzyme quality^1,3,9^. Varying digestion time impacts fluorescence intensity^1^ and how much the tissue expands in water, or the expansion factor^5^. To determine the effect of digestion time on the fluorescence intensity of the Amigo2-EGFP reporter line, which predominantly labels hippocampal area CA2 neurons, we performed a time course of enzymatic digestion with proteinase K as described in the proExM protocol^3^. Initially, we performed the time course on 40-micron vibratome cut sections from perfused adult Amigo2-EGFP mouse brains. Hydrogels were digested for 2, 4, 8 or 16 hours at room temperature. Unfortunately, there was insufficient EGFP signal remaining at the 8 and 16 hour timepoints to directly compare hydrogels across conditions (data not shown). We then repeated the experiment on sections immunostained for GFP prior to ExM and imaged the resulting gels with identical acquisition parameters to directly compare fluorescence intensities and expansion factors (Fig. 2). Importantly, we measured micro expansion factors, or the degree to which cell soma areas expanded versus the commonly reported macro expansion factors calculated by measuring how much the hydrogel expands. We found that average fluorescence intensities diminished as the length of digestion time increased (one-way ANOVA; F-stat: 20.96, p-value: 8.17E-14. N = two animals, 1-2 sections per animal per time point; 159 total cells, 32 ± 4 cells per time point, Fig. 2AB). Further, we found that average expansion factors increased as length of digestion time increased (one-way ANOVA; F-stat: 21.07, p-value: 4.26E-11, Fig. 2CD), resulting in an inverse relationship between fluorescence intensity and expansion factor (Fig. 2E). Interestingly, we did not detect significant decreases in fluorescence intensities from pairwise comparisons between the 8-hr digestion and the 2- or 4-hr digestions (p=0.90 and 0.55, respectively, Tukey’s post hoc test, Fig. 2D), despite significant increases in expansion factors between the same comparisons (p=0.001 for each). These data indicate that 8-hr digestion retains the most fluorescence for a sizable expansion factor (~3) that is not significantly different from the overnight expansion factor (p=0.18). However, we note that the fluorescence intensities reported here are corrected for background, and shorter digestion times have greater fluorescence background signal compared with longer digestion times (see Table 1). We also compared the cell area micro expansion factor to the extent that the tissue section expands (macro expansion factor) and found the macro expansion factor to be consistently greater than the micro expansion factor (Fig. 2F). Regardless of digestion time, we were able to successfully acquire robust fluorescent images at each time point, including overnight (Fig. 2G), as long as immunostaining was performed prior to ExM.

**Figure 2:**
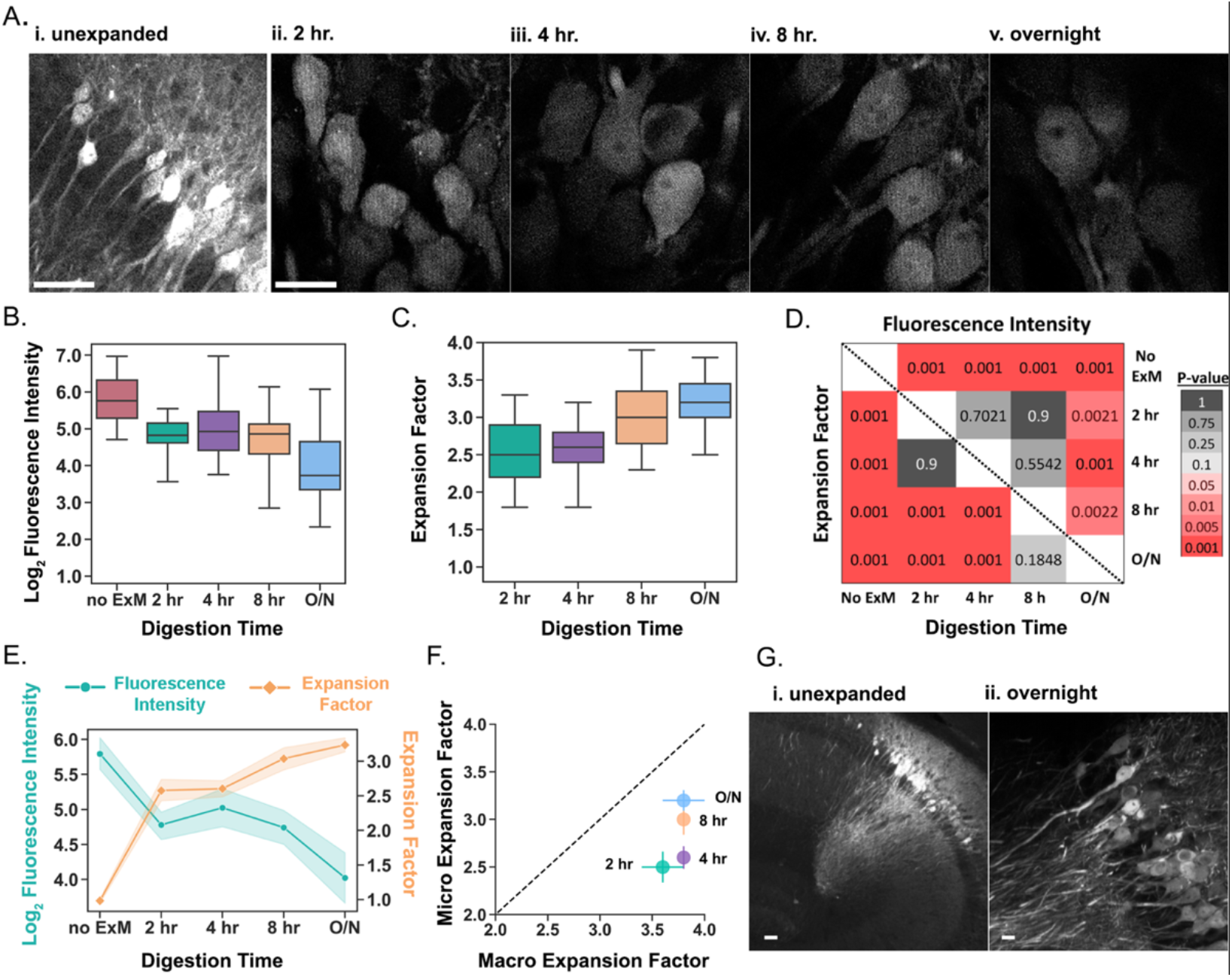
The effect of proteinase K digestion time on fluorescence intensity and expansion factor. **A)** Representative single z-section images of unexpanded (i) and expanded GFP+ CA2 neurons after 2-hour (ii), 4-hour (iii), 8-hour (iv), or overnight (v) digestion with proteinase K. Image acquisition parameters were identical for each condition, except for the unexpanded condition. **B)** GFP fluorescence intensity measured in the cell soma after the different digestion times. Fluorescence was corrected for background and log_2_ transformed. Line represents the median. Digestion time had a significant effect on GFP fluorescence (one-way ANOVA; F-stat: 20.96, p-value: 8.17E-14, 32 ± 4 cells per time point from two animals, 159 total cells). **C)** Expansion factor of the expanded cell somas in B. Expansion factors were calculated using cell soma areas relative to the average unexpanded cell soma area. Digestion time had a significant effect on expansion factor (one-way ANOVA; F-stat: 21.07, p-value: 4.26E-11). **D)** Matrix of p-values visualizing the results of pairwise Tukey’s post hoc tests comparing mean fluorescence (upper diagonal) or expansion factor (lower diagonal) at each digestion time point. Matrix is color coded by p-value (red = significant; grey = not significant; α=0.05). **E)** Effect of digestion time on fluorescence intensity (cyan; left axis) and expansion factor (orange; right axis). Plot shows the mean and 95% confidence intervals from B and **C.** **F)** Plot comparing cell area micro expansion factors versus tissue macro expansion factors. Error bars = 2*sem. **G)** Unexpanded (i) and overnight expanded (ii) max-projection 10X confocal images of GFP+ CA2 cells. Images are the same as in A(i) and (v). Imaging parameters were optimized separately to obtain the best image of both. Scale bars = **(A, G)** 50 μm.

**Table 1:** Digestion time course. Table showing mean expansion factor and fluorescence intensities (FI) for each digestion time point and the unexpanded control. Also included is the average background fluorescence intensity (“BG FI”), the normalized log2 transformed fluorescence, and the number of cells (“N Cells”) for each digestion condition.

| Digest Time | Exp Factor | Mean FI | BG FI | Log<sub>2</sub> Norm FI | N Cells |
| --- | --- | --- | --- | --- | --- |
| no ExM | 1.0 (±0.02) | 69.0 (±5.39) | 7.24 (±0.63) | 5.795 (±0.12) | 31 |
| 2 hour | 2.6 (±0.08) | 32.91 (±1.93) | 3.56 (±0.4) | 4.78 (±0.10) | 29 |
| 4 hour | 2.6 (±0.06) | 42.79 (±4.77) | 4.21 (±0.24) | 5.024 (±0.14) | 33 |
| 8 hour | 3.0 (±0.08) | 32.43 (±2.43) | 2.34 (±0.01) | 4.742 (±0.13) | 35 |
| overnight | 3.2 (±0.05) | 22.84 (±2.72) | 2.28 (±0.03) | 4.025 (±0.18) | 31 |

### Cells expand equivalently, independent of section depth

Next we considered if the time-dependent effect of proteinase K on expansion factor and fluorescence intensity is impacted by tissue depth. We reasoned that cells near the surface may have greater access to proteinase K and/or fluorescently-labeled antibodies compared with cells deeper within tissue that may impact expansion factor and/or fluorescence intensity, respectively. To test this, we compared cell soma areas binned by Z-section depth (adjusted by hydrogel thickness) across digestion time points.

We did not detect a systematic difference in cell soma area across z-section depth (Fig. 3), indicating that proteinase K equivalently digests 40-micron thick tissue, at least by the 2-hr time point. For some time points (e.g. 4- and 8-hrs) fluorescence intensity appeared brighter near the surface compared with deeper sections (Fig. 3B). However, we note this is likely due to optical limits, as fluorescence intensity was not brighter at the far-end surface that has equivalent access to proteinase K and antibody solutions. We saw similar results with 100-micron thick tissue (data not shown).

**Figure 3:**
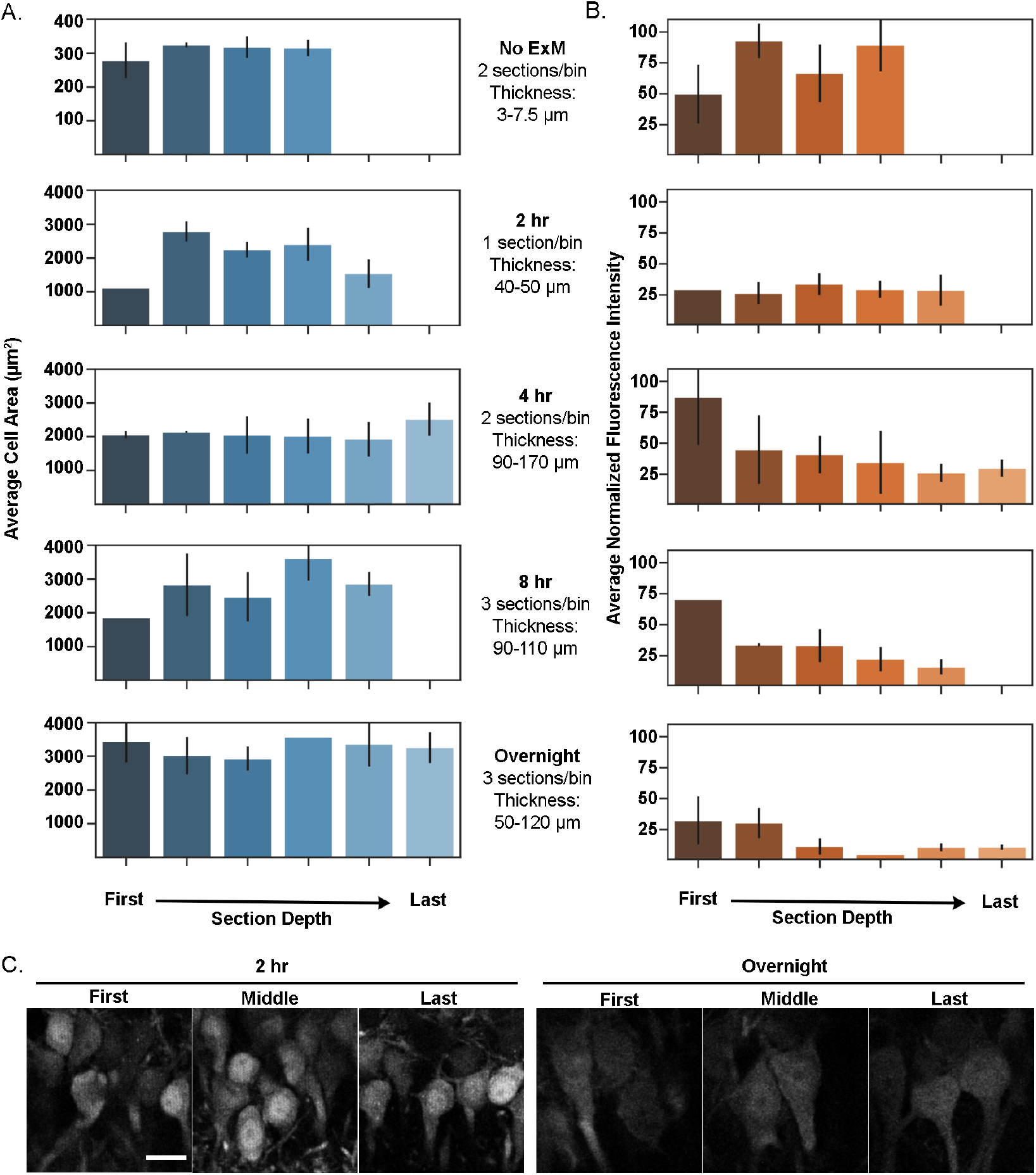
No appreciable effect of tissue depth on expansion factor. **A)** Average cell soma area (in μm^2^) by Z section depth for unexpanded control and digestion conditions. Cell soma area was measured in the Z section with the widest soma diameter. Because of the overall increase in depth with expansion, cells were binned by Z section to achieve ~6 bins per time point. N = 32 ± 4 cells per time point. **B)** Background subtracted average cell soma fluorescence by Z section depth. Binning of the data is the same as in A. The data are not log_2_ transformed as in Fig. 2. **C)** Representative confocal images of GFP+ CA2 cell somas after 2-hour (left panel) or overnight (right panel) digestion at three Z-section depths. Cell soma size and fluorescence is variable in the Z dimension in unexpanded conditions, which does not appreciably change under difference digestion conditions. Error bars show the standard deviation. Scale bar is 50μm.

### Immunolabeled organelles are best resolved with IHC prior to ExM

Protease digestion can decrease fluorescence intensity by impacting fluorescently-labeled antibodies and/or target antigen availability. Thus, immunohistochemistry (IHC) labeling before or after ExM (further referred to as IHC-pre and IHC-post, respectively) can be affected, but it is unknown if they equally affect the fluorescence intensity of antibodies that require antigen retrieval. Protease-free digestion protocols (e.g. detergent plus heat created in an autoclave liquid cycle) have been shown to effectively digest hydrogels, and avoids protease-dependent depletion of fluorescence intensity^3^. However, these protocols have not been tested with antibodies that require antigen retrieval. In Figure 4, we directly compared COX4-mitochondria labeling after protease (proteinase K) or autoclave (i.e. mild digestion) digestion ExM protocols^3^. We further compared these digestion protocols with IHC performed pre (Fig. 4AB) or post ExM (Fig. 4CD). Our optimized protocol for COX4 immunostaining on unexpanded sections (Fig. 4E) requires antigen retrieval (boiling) prior to IHC, thus an antigen retrieval step was included for sections bound for proteinase K digestion.

**Figure 4:**
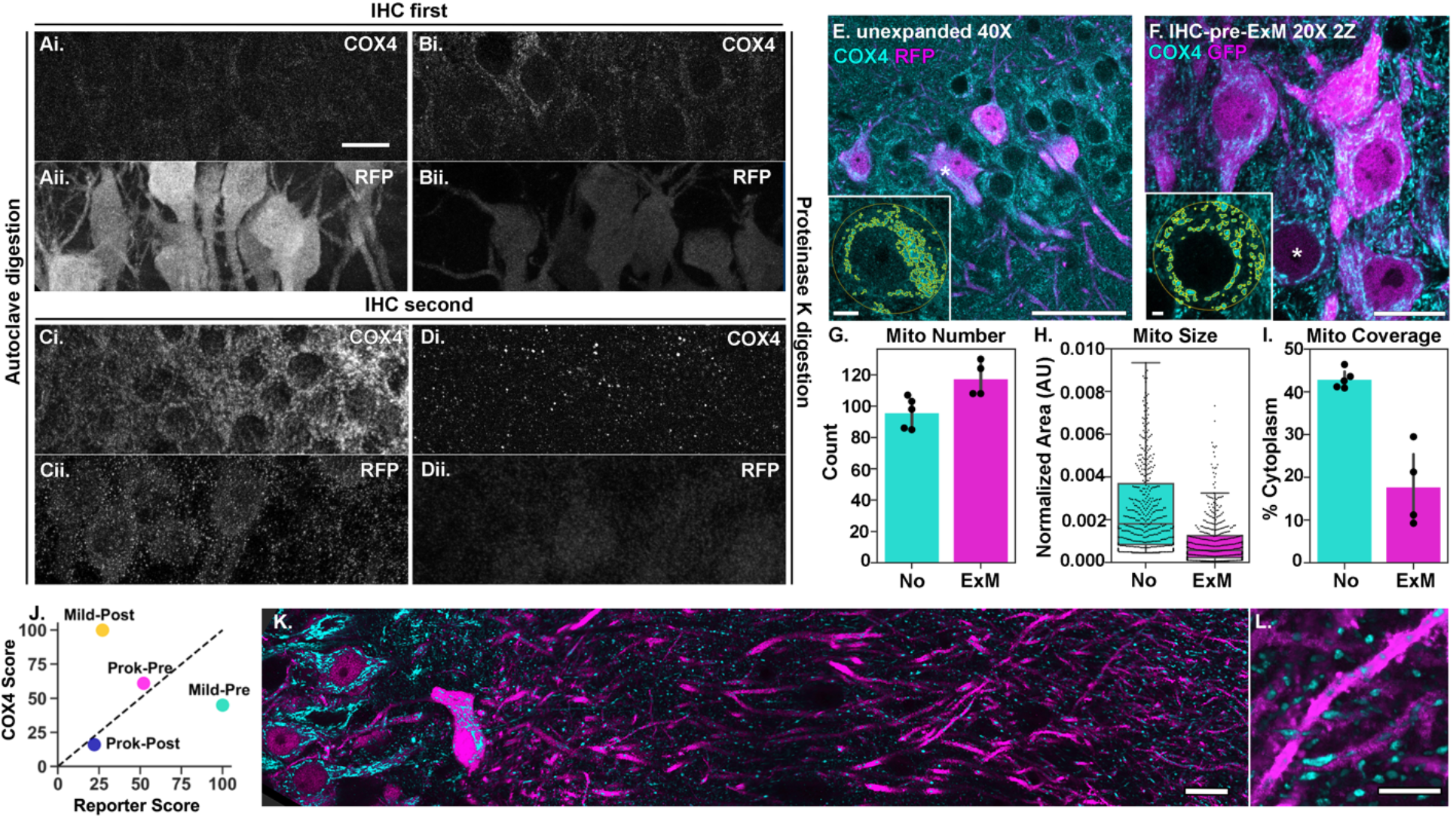
ProExM pipeline comparison for visualizing mitochondria. **A)** 20× Max-projection image of a section immunostained for COX4 (i) and RFP (iii) pre-ExM with autoclave digestion. Images in panels A-D were acquired with identical imaging parameters to directly compare conditions. Brightness and contrast were adjusted to the same extent across the entire image. **B)** 20X Max-projection image of a boiled section immunostained for COX4 (i) and RFP (ii) pre-ExM with overnight proteinase K digestion. **C)** 20X Max-projection image of a section immunostained for COX4 (i) and RFP (ii) post-ExM with autoclave digestion. **D)** 20X Max-projection image of a boiled section immunostained for COX4 (i) and RFP (ii) post-ExM with overnight proteinase K digestion **E)** 40X Max-projection image from a subset of Z sections from a boiled, unexpanded section immunostained for COX4 and RFP. Inset is a single Z section from the indicated representative cell (white asterisk) with mitochondria outlined as quantified in G-I. Imaging parameters were optimized per condition in **E-F.** **F)** 20X Max-projection image from a subset of Z sections from a boiled section immunostained for COX4 and GFP pre-ExM with overnight proteinase K digestion as in B. **G)** The average number of mitochondria per cell in unexpanded (E) and expanded (F) sections. **H)** Mitochondria area per cell (normalized by soma area). Line reflects median and whiskers are twice the inner quartile range. **I)** The average percent cytoplasm area (soma area - nuclear area) containing mitochondria. Note that ExM better resolves densely packed mitochondria, resulting in greater number, smaller sized, and fewer percent area per cell soma compared with no-ExM. **J)** Semi quantitative histological scores comparing tested conditions in panels A-D. **K-L)** Tile scan and high magnification image of mitochondria (cyan) in CA2 neurons and distal dendrites (magenta) with the conditions in **F.** Scale bars: 50 μm **(A,E,F,K),** 20 μm **(L)** and 5 μm (insets).

We detected COX4-labeled mitochondria using both proteinase K and autoclave digestion ExM protocols (Fig. 4), however proteinase K digested gels performed better with IHC-pre, and autoclave digested gels performed better with IHC-post. Compared with unexpanded COX4-labeled mitochondria, expanded COX4-labeled mitochondria were on average slightly greater in number per cell (ExM: 117.5 ± 5.6, No ExM: 95.8 ± 4.4) and smaller in size (mitochondria size: ExM 0.001 ± 0.0, No ExM 0.003 ± 0.0) after normalizing to cell soma area (total mitochondria area: ExM 422.5 ± 93.7, No ExM 123.8 ± 2.4) (Fig. 4G). To account for potential anisotropic expansion of cell somas, we compared the ratio of nuclear area (measured via DAPI) to the cell soma area (measured via reporter labeling) and found them to be similar on average with and without ExM (ratio: ExM 0.357 ± 0.00, No ExM 0.347 ± 0.01; nuclear area: ExM 1,383.1 ± 93.5, No ExM 154.5 ± 7.8; soma area: ExM 3,869.5 ± 259.2, No ExM 443.3 ± 11.8), indicating that the decrease in percent cytoplasmic area of mitochondria (Fig. 4I) is due to better individually resolved expanded mitochondria.

In regards to reporter labeling, RFP fluorescence only fared well when IHC was done prior to ExM, regardless of digestion method. Thus, for epitopes that require antigen retrieval and/or reporter labeling, IHC-pre-ExM is the preferred method of choice. Note that antigen retrieval diminishes reporter labeling (RFP and GFP) and explains the difference in RFP intensity between autoclave (Fig. 4Aii) and proteinase K (Fig. 4Bii) digested gels. If only mitochondria labeling is required, the autoclave digestion protocol performs well with IHC-post. The inferior staining of COX4 with IHC-pre with autoclave digestion compared with IHC-pre with proteinase K digestion may be due to a lack of antigen retrieval in the former that is achieved by autoclave digestion. To provide a semi-quantitative measure for each method tested in Fig. 4A-D, we scored the COX4 and reporter images on a scale from 0-100 based equally on the brightness of the signal and the quality of the labeling, the latter of which took into account the amount of noise and how closely the labeling pattern matched the expected pattern from unexpanded samples. A higher score reflects greater fluorescence signal or better signal quality as illustrated in Fig. 4J. The ProK-pre condition best preserves both mitochondria and the reporter labels. This can be seen in the spinning disk confocal images in Fig. 4K-L. With this combination, we can observe individual mitochondria within reporter-labeled dendrites (Fig. 4L).

We next compared the proteinase K and autoclave ExM protocols with GOLGA5-immunolabeling of Golgi apparatus (Fig. 5), which does not require antigen retrieval. Consistent with our COX4 results, GOLGA5-labeled Golgi were detected with either proteinase K or autoclave digested protocols. Golgi in expanded sections were well resolved and revealed complex Golgi structure compared with Golgi in unexpanded sections (Fig. 5F). In contrast to our COX4 results, GOLGA5-labeling fared well in both IHC-pre-ExM digestion conditions, likely due to robust GOLGA5-labeling in unboiled sections. Similar to COX4, GOLGA5-labeling post-ExM with proteinase K digestion was unsuccessful. IHC-pre with autoclave digestion was not tested. As with RFP reporter labeling, GFP reporter labeling also fared well with IHC-pre and autoclave digestion, albeit at lower intensities than proteinase K digestion. Thus, GFP and RFP fluorescence retention are comparable when antibody-labeled prior to ExM and they retain more fluorescence with proteinase K digestion compared with autoclave digestion. It is important to note that we detect qualitative differences in the ability of different reporter antibodies to detect spines, with and without ExM (see Table 3). Using the Prok-pre conditions, we were able to determine that there is no GOLGA5 labeled Golgi localized CA2 dendrites (Fig. 5G), despite clear GOLGA5 labeled Golgi in CA2 cell bodies (Fig. 5G-H). The GOLGA5 signal and reporter signal were semi-quantitatively scored as described above for mitochondria (Fig. 5I).

**Figure 5:**
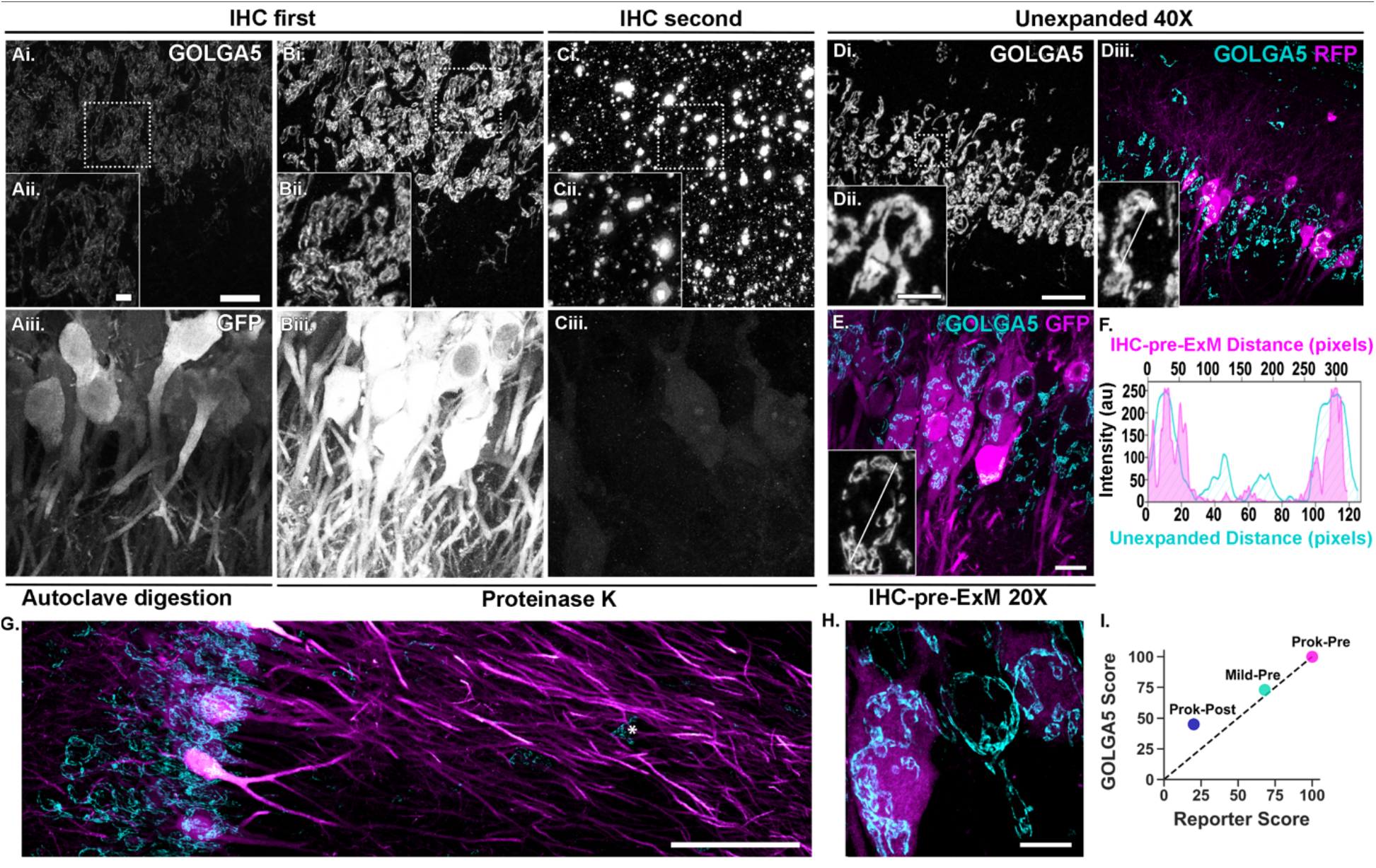
ProEXM pipeline comparison for visualizing Golgi. **A)** 20X Max-projection image of a section immunostained for GOLGA5 (i) and GFP (iii) pre-ExM with autoclave digestion. (ii) Insets are zoomed images of the dotted box in (i). Images in panels A-C were acquired with identical imaging parameters to directly compare conditions. Brightness and contrast were adjusted to the same extent across the entire image. **B)** 20X Max-projection image of a section immunostained for GOLGA5 (i) and GFP (iii) pre-ExM with overnight proteinase K digestion. **C)** 20X Max-projection image of a section immunostained for GOLGA5 (i) and GFP (iii) post-ExM with overnight proteinase K digestion. **D)** 40× Max-projection of a subset of Z sections from an unexpanded section immunostained for GOLGA5 (i) and RFP (iii). Imaging parameters were optimized per condition in D-E. **E)** 20X Max-projection of a subset of Z sections from image in **(B)** immunostained pre-ExM with overnight proteinase K digestion. Note the increase in Golgi complexity that is resolved with autoclave or proteinase K digested gels compared with unexpanded sections, although fluorescence intensity was better retained with proteinase K digestion. **F)** Line plot profile showing the fluorescence intensity of unexpanded Golgi (cyan) and Golgi after IHC-pre-ExM with proteinase K digestion (magenta). Measured Golgi are inset in Dili and **E.** **G-H)** Tile scan and high magnification image of GOLGA5 staining in CA2 neurons with the conditions in E. Note the lack of GOLGA5 signal in CA2 dendrites except that from glial cells (asterisk). **I)** Semi quantitative histological scores comparing tested conditions in panels A-C. Scale bars: 50 μm **(Ai, Di, E),10** μm **(Aii, Dii),** 200 μm **(G)** and 25 μm **(H).**

**Table 2:** Antibodies and conditions. Table showing primary and secondary antibody conditions used for expansion microscopy.

| Primary Antibody | Host | Clonality | Supplier | Catalogue # | Antigen retrieval | Concentration | Incubation |
| --- | --- | --- | --- | --- | --- | --- | --- |
| GFP | Chicken | Polyclonal IgY | Abcam | ab13970 | Not Required | 1:500 | 72+ hrs at RT |
| GFP Booster Atto 488 | Alpaca | Monoclonal VHH | Chromotek | gba488-100 | Not Required | 1:200 | 72+ hrs at RT |
| RFP | Rabbit | Polyclonal IgG | Rockland | 600-401-379 | Not Required | 1:500 | 72+ hrs at RT |
| RFP | Chicken | Polyclonal IgY | Rockland | 600-901-379 | Not Required | 1:250 | 72+hrs at RT |
| GOLGA5/Golgin-84 | Rabbit | Polyclonal IgG | Abcam | ab224040 | Not Required | 1:500 | 72+ hrs at RT |
| COX4 | Rabbit | Polyclonal IgG | Synaptic Systems | 298 003 | 5 min at 100C | 1:500 | 72+hrs at RT |
| ERGIC-53/P58 | Rabbit | Polyclonal IgG | Sigma Aldrich | E1031 | Citrate at 80C | 1:600 | 72+ hrs at RT |
| ST3GAL5 | Rabbit | Polyclonal IgG | Sigma Aldrich | AV46358 | Citrate at 80C | 1:250 | 72+ hrs at RT |
| MCU | Rabbit | Polyclonal IgG | Sigma Aldrich | HPA016480 | 5 min at 100C | 1:500 | 72+ hrs at RT |

| Secondary Antibody |  |  |  |  |  |  |  |
| --- | --- | --- | --- | --- | --- | --- | --- |
| Goat anti Mouse IgG (H+L) Alexa 488 | Goat | Polyclonal IgG | Invitrogen | A11029 | NA | 1:500 | 48+ hrs at RT |
| Goat anti Mouse IgG (H+L) Alexa 546 | Goat | Polyclonal IgG | Invitrogen | A11030 | NA | 1:500 | 48+ hrs at RT |
| Goat anti Rabbit IgG (H+L) Alexa 488 | Goat | Polyclonal IgG | Invitrogen | A11034 | NA | 1:500 | 48+ hrs at RT |
| Goat anti Rabbit IgG (H+L) Alexa 546 | Goat | Polyclonal IgG | Invitrogen | A11035 | NA | 1:500 | 48+ hrs at RT |
| Goat anti Chicken IgY (H+L) Alexa 488 | Goat | Polyclonal IgY | Invitrogen | A11039 | NA | 1:500 | 48+ hrs at RT |
| Goat anti Chicken IgY (H+L) Alexa 546 | Goat | Polyclonal IgY | Invitrogen | A11040 | NA | 1:500 | 48+ hrs at RT |

**Table 3:** Detection of reporter labeled spines

| Tissue/Antibody | ExM (protK) | IHC (pre-ExM) | Spines |
| --- | --- | --- | --- |
| Am2-GFP | unexpanded | no IHC | + |
|  |  | ag-no IHC | - |
|  | expanded | no IHC | - |
|  |  | ag-no IHC | - |
| Am2-GFP/GFP-chicken | unexpanded | IHC | ++ |
|  |  | ag-IHC | + |
|  | expanded | IHC | +++ |
|  |  | ag-IHC | ++ |
| Am2-GFP/GFP booster-alpaca | unexpanded | IHC | + |
|  |  | ag-IHC | - |
|  | expanded | IHC | - |
|  |  | ag-IHC | nt |
| Am2-icre;tdTomato | unexpanded | no IHC | + |
|  |  | ag-no IHC | + |
|  | expanded | no IHC | - |
|  |  | ag-no IHC | nt |
| Am2-icre;tdTomato/RFP-chicken | unexpanded | IHC | +++ |
|  |  | ag-IHC | + |
|  | expanded | IHC | ++ |
|  |  | ag-IHC | + |
| Am2-icre;tdTomato/RFP-rabbit | unexpanded | IHC | ++++ |
|  |  | ag-IHC | nt |
|  | expanded | IHC | ++++ |
|  |  | ag-IHC | nt |
| - not detected; + rare SLM only; ++ OK; +++ Good; ++++ Excellent; nt Not tested |  |  |  |

### Subcellular structures in the tissue are minimally distorted after expansion with proExM

To confirm the IHC-pre-ExM protocol with 8 hours proteinase K digestion reliably maintains macro and micro tissue structure, we measured the macro expansion of the whole tissue section (Fig. 6AB) and the micro expansion of individual cells for 5 different animals (Fig. 6C). The average macro expansion factor was 3.88 and the average micro expansion was 3.33. Example tile images of the same tissue before and after expansion are shown in Fig. 6DE. To quantify the amount of distortion, we performed a root mean squares (RMS) analysis on three sets of GOLGA5 images of the same field of view before and after proExM, as described by Chozinski et. al^1^. The post-ExM image (Fig. 6H) was registered to the preExM image (Fig. 6G) in two steps, with a rigid and then a non-rigid B-spline registration in Elastix. A vector field map was generated (Fig. 6I) and RMS was calculated with code provided by Chozinski et. al. (see Methods). Over a length of 10 μm, the average RMS error across the three animals was 0.2 μm, which is a 2% error. This is in line with previous publications^10^ and demonstrates little distortion between the pre-ExM images and the post-ExM images.

**Figure 6:**
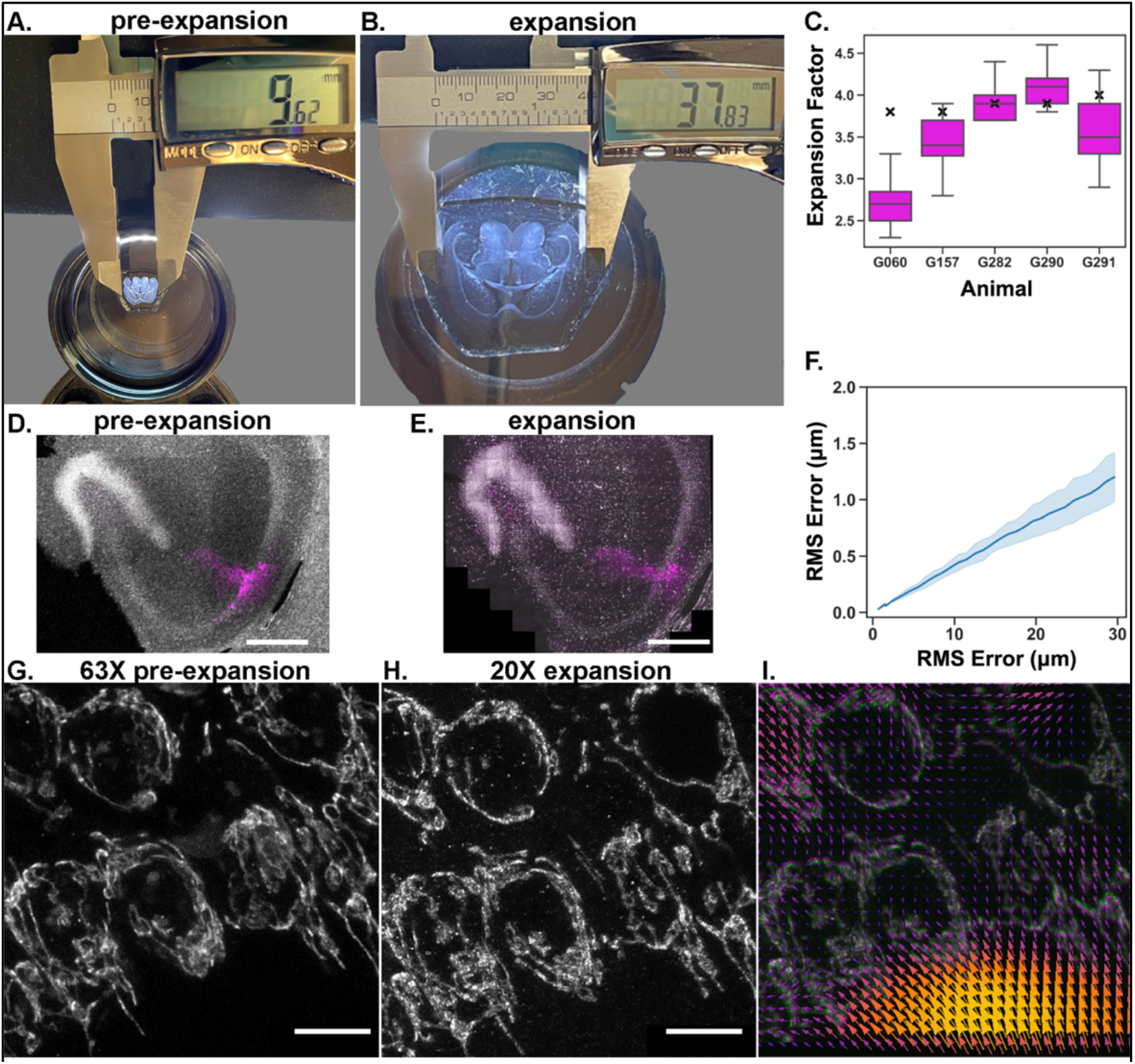
Minimal distortion of tissue or subcellular structures after ProExM. **A-B)** Representative hydrogels from pre-expanded **(A)** and expanded **(B)** horizontal brain sections after 8-hour digestion with proteinase K. C) Micro and macro expansion factors from 5 mice. Box plots show the calculated cell body micro expansion factors (see methods) and X denotes the macro expansion factor of the tissue for each hydrogel as measured in B. **D-E)** Representative confocal images of DAPI positive nuclei (grey) and EGFP positive CA2 neurons (magenta) in pre-expanded (5X) and expanded (10X) hippocampus after 8-hour digestion with proteinase K. F) Root mean square (RMS) error plot calculated from N=3 gels from 3 mice. **G-l)** Representative confocal images of GOLGA5 (grey) immunostaining preExM in 63X unexpanded **(G)** and 20X expanded (H) CA2 neurons after 8-hour digestion with proteinase K. The resulting vector plot (I) from B-spline image registration. Arrows indicate the direction and magnitude of the transformation required to align the expanded image to the pre-expanded image.

### Dendritic spines are best resolved in expanded GFP or RFP-immunolabeled tissue

In addition to resolving subcellular organelles such as mitochondria and Golgi apparatus, the ability to resolve fine dendritic structures such as spines allows us to address questions about function and plasticity at synapses. Thus, we set out to find the optimal combination of ExM conditions for subcellular organelles and reporter antibodies to resolve dendritic spines in EGFP and tdTomato reporter mice. Table 3 compares spines in unexpanded and expanded tissue, with or without immunolabeling for the reporter protein, and with or without antigen retrieval by boiling or citrate. Expanded samples were immunostained pre-ExM (if at all) and digested overnight with proteinase K, as described in the methods. To boost the signal of the reporter protein, we tested two antibodies against each EGFP and tdTomato reporters (anti-GFP and anti-RFP, respectively). The ability to resolve dendritic spines in each condition was qualitatively assessed by multiple investigators (not blinded to condition), and indicated by the number of + in the table (from + to ++++). A greater number of + indicates better discrimination of spines and indicates dendritic spines could not be resolved for a given condition. Some conditions in the table have yet to be tested as indicated where applicable.

Figure 7 shows representative images of dendritic spines for each of the four conditions tested. We saw the best resolution of spines in expanded tdTomato+ tissue that was stained with the rabbit anti-RFP without any antigen retrieval (Fig. 7A; “Am2-icre;tdTomato/RFP-rabbit” in Table 3). We noted that the chicken RFP antibody did not label spines as well as the rabbit RFP antibody (“Am2-icre;tdTomato/RFP-chicken”, Table 3), indicating that not all reporter antibodies are equal when it comes to labeling spines. In general, spines were easier to resolve with proExM in tdTomato reporter mice (top panel) compared to EGFP reporter mice (bottom panel). The reason is likely multifactorial: a combination of a better performing RFP antibody, a difference in fluorescence retention between tdTomato and EGFP after proExM^2^, and a difference in fluorescence retention between their secondary antibodies (Alexa546 and Alexa488, respectively) after proExM^2^.We find CA2 spines are difficult to visualize in either mouse strain without prior immunolabeling. Thus, we do not believe mouse strain differences account for the differences in spine labeling after proExM. Compared to unexpanded tissue with the same IHC conditions, the proExM protocol increases the resolution of spines by increasing their physical size and separation from nearby dendritic branches and reducing background fluorescence and/or light scattering^11^ to enable spine morphometric analyses on individually traced neurons.

**Figure 7:**
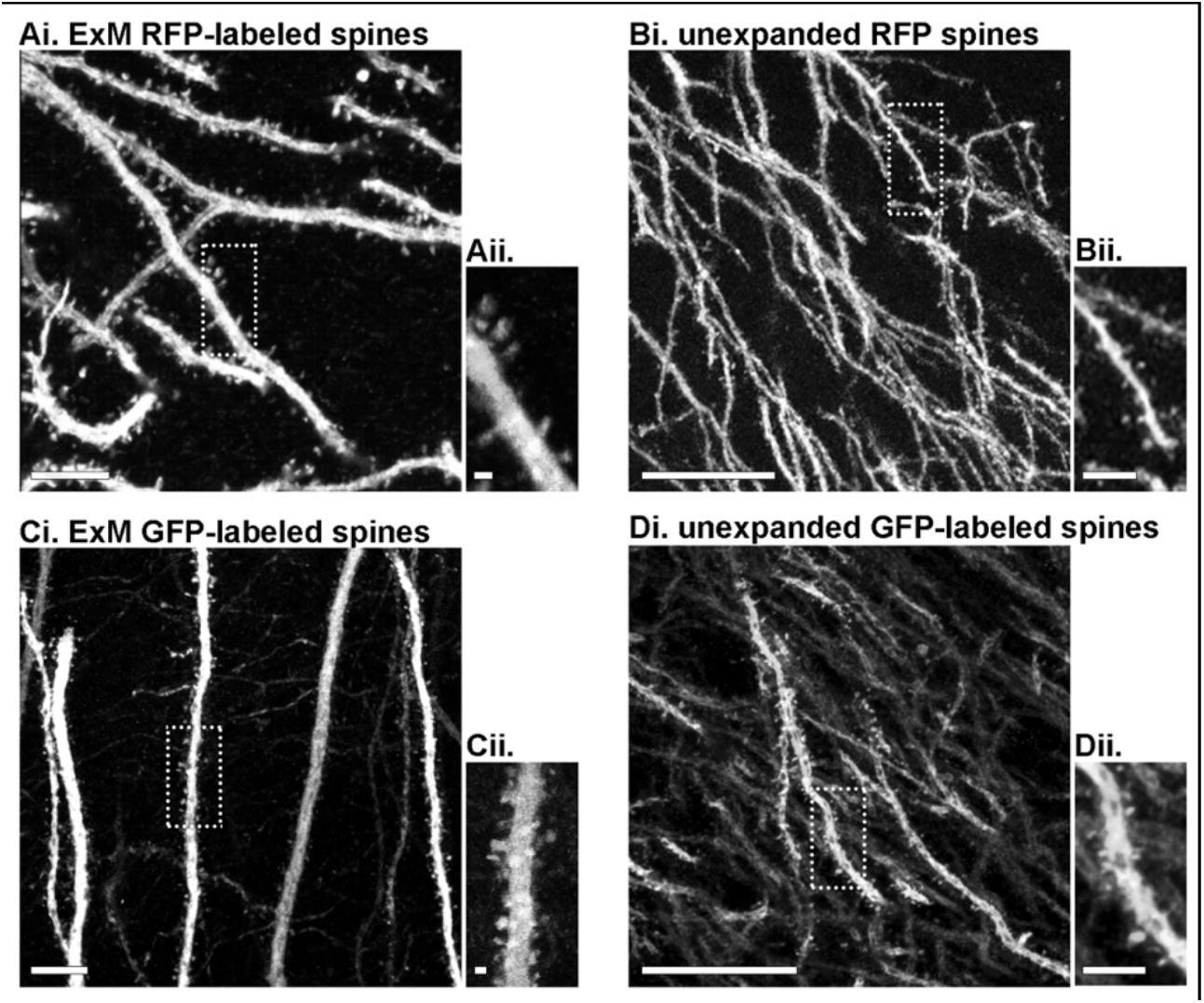
Resolving dendritic spines with ExM. **A)** 20X with 3X zoom max-projection image (i) of a section immunostained for RFP pre-ExM with overnight proteinase K digestion. This mouse received 2 tamoxifen injections. (ii) Insets are 2X zoomed images of the dotted box in (i). Images were acquired with parameters optimal per condition. Brightness and contrast were adjusted per condition across the entire image. **B)** 40X with 2X zoom max-projection image of an unexpanded tdTomato section. This mouse received 2 tamoxifen injections. **C)** 20X with 2X zoom max-projection image of a section immunostained for GFP pre-ExM with overnight proteinase K digestion. **D)** 40X with 2X zoom max-projection image from an unexpanded section immunostained for GFP. Note the increased background labeling and density of dendritic branches in the unexpanded images, making dendrites more difficult to trace compared with expanded images. Scale bars: 25μm **(Ai, Bi, Ci. Di)** and 5μm **(Aii, Bii, Cii, Dii).**

## DISCUSSION

ProExM is a powerful tool that increases the resolution of conventional fluorescence microscopy to ~70 nm and can be performed with tools available in a standard molecular biology laboratory^10,12^. Because a fully expanded hydrogel is mostly water, the optically clear sample is well suited to resolve densely packed organelles and tissues. However, the digestion and expansion process greatly reduces fluorescence retention making it necessary to optimize ExM conditions per sample for specific end goals. Here we described a proExM workflow optimized for resolving subcellular organelles (mitochondria and the Golgi apparatus) and spines in fixed mouse brain tissue. We reliably found that immunostaining before proExM (IHC-pre-ExM) and using a proteinase K based digestion for 8 hours resulted in the best fluorescence signal to resolve subcellular organelles while maintaining sufficient reporter labeling to visualize spines and trace individual neurons. With these methods, we were able to more accurately quantify mitochondria size and number and better visualize Golgi ultrastructure in reconstructed CA2 cell bodies in the hippocampus.

Several groups have optimized expansion protocols to visualize subcellular organelles across different sample types, including various cell lines^1,5,8,13–16^, rat liver^17^, clinical specimens^18^, fungi^6,19^, songbird^20^ and drosophila^21,22^. Others have used ExM to visualize subcellular structures, including mitochondria^1,2,23–25^ and/or spines^2,20,23^ in brain tissue, but few have systematically analyzed how fluorescence intensities and expansion factors compare across protocols or with unexpanded measurements. This is critically important as several groups have noted discrepancies in micro versus macro expansion factors in other sample types^7,14,17,26–28^, including dissimilarities in expansion factors of different subcellular organelles^14^ or of subcellular organelles across neighboring cells and tissues^17^. Others, however, have reported minimal to no differences in micro vs macro expansion factors^6,19^. In our measurements, we found discrepancies between the micro and macro-expansion factors. On average, the 8-hour proteinase K digestion produced a micro expansion factor of 3.33 and a macro expansion factor of 3.88 (Fig. 6C). Interestingly, the macro expansion factor was relatively insensitive to digestion time past 2 hours, while the micro factor continued to increase (Fig. 1F). Our average micro expansion factors are lower than the commonly reported 4-4.5X macro expansion factor for proExM, which is consistent with other reports using micro expansion factor measurements^14,15^ (but see also ref^26^), reinforcing the notion that each sample type needs to be independently optimized and validated for ExM.

To determine the optimal digestion time for fluorescence retention in fixed brain sections, we performed a digestion time course and found the greatest drop in fluorescence after the overnight digestion. There was no significant drop in fluorescence between 2, 4 or 8 hours of digestion. However, there was a significant increase in expansion factor during this time period. Expansion factor begins to plateau after 8 hours of digestion, and while there is a slight increase in expansion factor in the overnight condition it is not statistically significant. Therefore, under the conditions used here, a digestion time of 8 hours is ideal for achieving a robust expansion (~3X) without further loss of fluorescence seen with overnight digestion. At all digestion time points, the expansion factor did not systematically vary by depth, indicating that 2 hours in proteinase K is sufficient for uniform penetration and isotropic expansion in the Z dimension of 40 micron brain sections, as previously reported for thicker sections and longer digestion times^10^.

In regards to labeling subcellular organelles in fixed brain sections, we were able to visualize expanded mitochondria with a COX4 antibody and expanded Golgi apparatus with a GOLGA5 antibody, using either the proteinase K digestion or the mild autoclave digestion. In our hands, the IHC-pre ExM with proteinase K digestion outperformed the other conditions based on fluorescence signal retention for both immunostained organelles and reporter proteins. While EGFP and tdTomato have been reported to have different percent fluorescence retention after ExM^2^, they perform comparably when antibody-labeled prior to ExM, as recently reported in cultured cells^29^. They also retain more fluorescence with proteinase K digestion compared with autoclave digestion. However, if the goal is only to visualize mitochondria, the mild autoclave digestion with IHC-post ExM also produced good COX4 staining, as seen for other mitochondria immunostains, like TOMM20^2^. The GOLGA5 antibody performed decently using IHC-pre ExM and autoclave digestion, which the COX4 antibody did not, likely due to COX4 immunostaining requiring antigen retrieval. Neither antibody performed well with IHC-post ExM and proteinase K digestion. It is important to note that conducting immunostaining prior to proExM introduces small positional errors due to linking the fluorophores into the gel. Immunostaining targets with primary and secondary antibodies imposes a linkage error of ~17.5 nm^30,31^. This can cause a localization error between a protein of interest and its fluorophore in an expanded state, however, the relative distance of the fluorophore to the epitope stays unchanged. This is in contrast to post-ExM labeling that leads to a relative smaller antibody size^6^.

Despite diminished fluorescence, ExM afforded better resolution for quantification of subcellular organelles compared to unexpanded organelles. We quantified the number and size of expanded mitochondria and found that the expanded mitochondria were smaller and more numerous than unexpanded mitochondria. This presumably is indicative of tightly packed mitochondria in the unexpanded samples being lumped together that can be separately resolved with expansion. Here, we normalized subcellular measurements (i.e. mitochondria size) to within cell nuclear and cytoplasmic areas. Another study found discrepancies in cell soma vs nuclear expansion factors^17^, but here the ratio of nuclear area to cytoplasmic area remained constant between unexpanded and expanded states, indicating isotropic expansion, perhaps due to our longer digestion times.

The fine details of the Golgi cisternae were also better resolved after expansion, whereas without expansion the Golgi were smoothened and much of the details lost. For the goal of visualizing reporter-labeled dendritic spines, we found that the addition of IHC-pre ExM was necessary for the resolution of both EGFP+ and tdTomato+ spines. Dendritic spines were best resolved in IHC-pre ExM with proteinase K digestion with either GFP or RFP antibodies. The ability to label dendritic spines in ExM was antibody dependent, with some antibodies against the same reporter faring better than others under the same IHC conditions.

Our lab has begun applying the described proExM methods to answer open questions involving subcellular organelles in neurons. One such open question is whether there is Golgi present in dendrites, or if the Golgi is limited to neuronal cell bodies. It is known that local translation of RNA occurs in dendrites^32^; however, there is conflicting evidence of the existence of Golgi apparatus in the dendrites, which would normally process newly transcribed membrane bound proteins. Using combined GOLGA5-labeling of Golgi apparatus and reporter neuron labeling with the proExM protocol described here, we do not detect GOLGA5 staining outside the cell soma or very proximal dendrites, consistent with reports that canonical Golgi markers are retained in the soma and not present in distal dendrites^25,34,35^.

### Considerations for subcellular imaging of expanded samples

The benefits of expansion come at several costs, including diminished signal concentration, hydrogel mechanical integrity and movement, and increased imaging volume and time^5^. Here we comment on the proposed workarounds for these issues that have or have not worked well for subcellular imaging of expanded brain sections.

Expansion microscopy substantially increases the thickness of a sample, which limits its ability to be imaged with standard high magnification microscope objectives with limited working distances. It can be expected that the gel thickness will be equivalent to 4-fold the depth of the gelation chamber. Further, keeping the tissue in plane during gel chamber assembly is difficult, often resulting in increased sample z-distance. Minimizing tissue thickness (40 microns versus 100 microns) and using a single coverslip for the gelation chamber helped minimize gel thickness without sacrificing ability to reconstruct neurons. However, in our hands, digestion time need to be decreased to 4 hours when using a single coverslip versus 8 hours for two coverslips as optimized here.

Loss of fluorescence due to the digestion of antibodies or epitopes and dilution of fluorescent molecules per unit volume can result in low contrast samples not suitable for high resolution imaging even with overexpressed fluorescent reporter proteins. We found that performing IHC beforehand and limiting the proteinase K digestion to 8 hours largely negated this issue. When the fluorescence signal is insufficient, imaging in low concentrations of PBS (0.5X PBS instead of 0.0001X PBS or water) substantially improved the contrast by increasing the concentration of fluorescent molecules per area. This will decrease the expansion factor, but in our hands, even a 2-3-fold expansion in optically clear tissue produces better resolved images of subcellular structures than unexpanded tissue.

Hydrogel movement during imaging is another common issue. Because gels expand 4-fold in x, y and z compared with unexpanded brain sections, tile scans are required to reconstruct entire neurons even at 10x (see Fig. 2Fii for a 10x single field of view of expanded neurons). Tile scanning increases the length of time required to image a gel and thus worsens gel shift. Gel shift was most noticeable while imaging on an upright microscope equipped with a water immersion lens, since the gel can easily shift when submerged. Gel shift was less noticeable when imaging either on an inverted scope or on an upright scope with air objectives (if the gel was dried for 30 minutes to adhere to the glass bottom plate), but these options negatively impact objective working distance or resolution, respectively. To minimize gel shift during upright imaging with a water dipping lens, we applied the following techniques to stabilize the gel in the imaging chamber (50 mm WillCo Well). Following full expansion in water, we surrounded the gel with 2% agarose in the imaging chamber. This noticeably reduced gel movement during acquisition of single images but shift was still detected during longer tile scans. We also tested re-embedding the gelated sample in an unexpandable gel and covalently linking it to the glass imaging dish^2^. This completely eliminated movement of the gel and allowed us to take long tile scans on an upright or inverted microscope. Unexpectedly, however, this also seemed to dampen the fluorescence signal, which was irreversible.

## METHODS

### Animals

Adult male and female Amigo2-EGFP (RRID:MMRRC 033018-UCD, bred for at least 10 generations onto C57BL/6J background) or Amigo2-icreERT2;RosaTdTomato transgenic mice were used. Amigo2-icreERT2;RosaTdTomato mice were generated by crossing Amigo2-icreERT2 mice^36^ to Ai14 mice (Jax #007914). Amigo2-icreER;ROSA-TdTomato mice were given 2 or 3 daily intraperitoneal injections of tamoxifen (Sigma T5648, 100mg/kg freshly dissolved daily in 100% ethanol then diluted 10-fold in corn oil and heated at 60C for 1 hour until in solution). Mice were group-housed under a 12:12 light/dark cycle with access to food and water *ad libitum.* All procedures were approved by the Animal Care and Use Committee of Virginia Tech.

### Immunofluorescence

Mice were anesthetized with 150mg/kg fatal plus solution and perfused with ice-cold 4% paraformaldehyde. Amigo2-icreERT2;RosaTdTomato mice were perfused one week post tamoxifen injections. Brains were dissected and post-fixed for at least 24 hours before sectioning 40 μm thick sections in the horizontal plane on a vibratome (Leica VT1000S). Sections to be stained with COX4 underwent antigen retrieval by boiling free floating sections in 1.7ml tubes for 3 min in nanopure water. All sections were washed in PBS and blocked for at least 1 hour in 5% Normal Goat Serum (NGS)/0.3% Triton-100x. See Table 3 for primary and secondary antibodies and conditions. Antibodies were diluted in blocking solution and sections were incubated for 72+ hours at room temperature (RT). After several rinses in PBS-T (0.3% Triton-100x), sections were incubated in secondary antibodies for 48 hours at RT. Prior to imaging, unexpanded sections were washed in PBS-T and mounted under Vectashield fluorescence media with 4’,6-diamidino-2-phenylindole (DAPI) (Vector Laboratories).

### Protein expansion microscopy solution preparation

Solutions were prepared as described by Asano et. al 2018^3^. Anchoring stock solution was prepared by dissolving Acryloyl-X, SE (ThermoFisher A20770) in DMSO (1:100 w/v) and stored at −20C. Monomer solution components were prepared by dissolving Sodium Acrylate in npH20 (33% w/v, Sigma 408220), Acrylamide in npH20 (50% w/v, Sigma A9099), N, N’-Methylenebisacylamide in npH20 (2% w/v, Sigma M7279). Monomer working solution was prepared by adding 2.25mL of 33% SA solution (8.6% w/v), 0.5 mL of 50% Acrylamide solution (2.5% w/v), 0.75 mL of 2% N,N-Methylenebisacrylamide solution (0.15% w/v), 4 mL of 5M NaCl (11.7% w/v), and 1 mL of 10X PBS. Inhibitor stock was prepared by dissolving 4-Hydroxy-TEMPO (0.5% w/v, Sigma 176141) in npH2O and initiator stock was made by dissolving Ammonium persulfate in npH2O (10% w/v, Sigma 248614). Accelerator solution was prepared by diluting TEMED in npH2O (10% v/v, Sigma T7024) immediately before use. All solutions except the TEMED accelerator were prepared before use and stored at −20C.

### Protein expansion microscopy

4X protein expansion microscopy (proExM) was carried out on horizontal mouse brain sections containing dorsal hippocampus as described in Asano et al 2018^3^. Sections were incubated overnight in Acryloyl-X stock/PBS (1:100, ThermoFisher, A20770) at RT in the dark. Following incubation, the slices were washed twice with PBS for 15 minutes each at RT. The gelation solution was prepared by adding 384 uL of monomer solution, 8 uL 4-Hydroxy-TEMPO inhibitor (1:200 w/v, Sigma Aldrich, 176141), 8uL TEMED accelerator (10% v/v, Sigma Aldrich, T7024), and lastly 8uL of APS initiator (10% w/v, Sigma Aldrich, 248614) for each section. Sections were then incubated in the gelation solution for 30-45 minutes at 4C in the dark. Gelated sections were placed on gelation chambers constructed from microscope slides with coverslips as spacers. Our gelation chambers produce gels with the thickness of two type No. 1.5 coverslips (~0.3mm thick). The chambers were filled with gelation solution and allowed to incubate at 37 C for 2 hours in a humidified container. Following gelation incubation, the gelation chamber was deconstructed to uncover the gelated brain section. To remove the gel from the chamber, digestion solution without proteinase K was applied and a coverslip was used to gently remove the sample. Digestion solution containing proteinase K (8U/mL, New England BioLabs, P8107S) was applied to gels and allowed to digest for 2-16 hours (see Results) at room temperature. Upon completion of digestion, gels were stained with DAPI (Sigma Aldrich, D9542, 1:10,000 in PBS) for 10 minutes at room temperature with shaking. The gels were then washed twice for 10 minutes with npH2O to remove excess DAPI and fully expand the gel.

#### Stabilizing ExM Gels with Agarose

To prevent movement during imaging, gels were fully expanded in water or 0.001X PBS in WillCo wells (HBSB-5040) and reversibly immobilized by applying liquid 2% agarose around and on top of the gel (in areas not containing tissue). Following application, the gel embedded with agarose was placed at 4C for at least 15 minutes to allow the agarose to fully solidify prior to imaging.

#### Re-embedding and Linking Gels to Imaging Dish

Re-embedding and covalently linking gels to a WillCo well allowed long-term imaging without gel shifting as described by Tillberg et al 2016^2^. First, the gel was completely expanded in water and then incubated in a non-expanding re-embedding solution (3% w/v acrylamide, 0.15% w/v N,N-methylenebisacylamide, 0.05% w/v APS, 0.05% w/v TEMED, and 5mM Tris). The gels were incubated with shaking at room temperature for 20 minutes. The gel was then transferred to a Bind-Silane treated WillCo Well plate and covered with a coverslip. Fresh re-embedding solution was then lightly applied surrounding the sample, which was then incubated at 37C for 1.5-2hrs without shaking. Once the re-embedding solution gelated, the coverslip was removed from the covalently linked gel and could be imaged.

#### Bind Silane Treatment of Imaging Dishes

Immediately before use, imaging dishes were treated with a bind silane solution as described by Tillberg et al 2016^2^. Before bind silane treatment, imaging dishes were briefly washed with npH2O, 100% Ethanol, and then allowed to dry. Bind silane solution (80% v/v EtOH, 2% v/v acetic acid, 0.05% v/v Bind-Silane) was then applied with shaking for 5 minutes. The dish was then washed with 100% EtOH and allowed to dry before usage.

#### Digestion Time Course Experiment

To assess the effect of digestion time on tissue expansion and fluorescence retention, we ran a digestion time course experiment by varying the amount of time the ExM gels were in digestion solution-- either 2 hours, 4 hours, 8 hours or 16 hours (overnight). The brains of two Amigo2-EGFP mice (One male and one female, 21-23 weeks old) were processed as above. During a pilot run, we noted that samples with digestion > 2 hours lost the majority of EGFP fluorescence, making it impossible to acquire images with identical parameters for direct comparison. Therefore, to boost the EGFP signal, approximately ten sections per brain (two sections per condition per mouse) were first immunostained in a 24-well plate as described in the immunohistochemistry methods, with a primary chicken antibody against EGFP (Abcam, ab13970; 1:500 concentration) at RT for 3 days, followed by a secondary antibody (Invitrogen Alexa-488, A11039; 1:500 concentration) incubation for 48 hours. As a control, a few sections were set aside after immunostaining to mount on slides without expanding.

Sections processed for ExM were anchored in Acryloyl-X overnight, washed twice with 1X PBS and incubated in gelation solution for 30 min at 4C the following day (see “Protein Expansion Methods” for more detail). The gels were incubated in a humid environment at 37C for 2 hours to set, and then carefully removed from the chamber and placed in a digestion solution with 8 U/mL of proteinase K (New England Bio, Cat # P8107S) for the designated period of time. All gels in this experiment were digested on a shaker at room temperature. At the end of the digestion time, the digestion solution was replaced with 1X PBS several times to wash the gels. Gels were stored in 1X PBS in the dark at 4C until imaging, at which point the PBS was replaced with npH2O to fully expand the gels. Gels were imaged at 10X (EC-Plan-Neofluar lens; 10x; 0.3 NA; 5.2mm WD) taking 10*μ*m steps for distance on a Zeiss 710 confocal microscope with the same image acquisition parameters, as described in more detail under “image acquisition”.

#### Digestion Time Course Analysis

The ExM images from the time course experiment were analyzed with the image processing program Fiji (v.+03..30, NIH). Before analysis, images were flattened across the Z dimension with the *“ZProject”* function to get an idea of how many GFP positive cells were present. This flattened image was only used for the selection of cells. Cells were excluded from analysis if they met any of the following criteria: 1) Incomplete cell (i.e. on the image border); 2) Any cell overlapping in Z with a cell already analyzed; 3) Cells with an ambiguous border or that were difficult to differentiate. Given these exclusion criteria, an average of 32 cells were analyzed for each time point across both animals. For some time points, cells were included from multiple hippocampal sections from the same animal.

For the fluorescence analysis, each cell body meeting the inclusion criteria was manually traced with the freehand selection tool in a single z section at the widest point of the cell. Once the cells were traced, the *“measure “* tool in Fiji was used to measure the mean intensity and the area within each individual cell ROI. To subtract background signal, one cell ROI was selected (at random) and moved to a location without EGFP signal and the background mean intensity was measured. The background mean intensity value was subtracted from the mean fluorescence intensity of each cell from the same image, resulting in a normalized mean fluorescence. The area of the cell body ROIs were used to calculate expansion factor by comparing it to the average area of 37 unexpanded cell bodies, which was calculated to be 300 μm^2^. To calculate the linear expansion factor for each cell, we used the below formula:

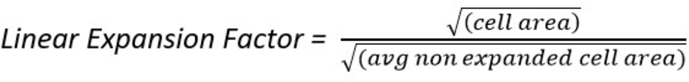

A total of 159 cells were included in the analyses (average of 32 +/− 4 cells per time point from two brains). The mean fluorescence and expansion factor measurements were imported into Python (v 3.7, installed with the Anaconda distribution), where the average mean fluorescence and average linear expansion factor were calculated for each time point, including the unexpanded control (see Fig. 2). The mean fluorescence data was not normally distributed (Shapiro-Wilk test for normality; p < 0.0001), so the mean fluorescence data was log2 transformed and 11 cell means were removed as outliers, pre-defined as two standard deviations from the mean. These same outliers were also removed from the expansion factor data; however, because this data was normally distributed (Shapiro-Wilk; p = 0.235), it was not transformed. One-way ANOVAs were run using an ordinary least squares model with the StatsModels package (ols, ANOVA_lm, statsmodels v0.11.1) comparing fluorescence intensity and linear expansion factors across digestion time points. After overall significance was reached, pairwise post-hoc Tukey’s multiple comparison tests (pairwise_tukeyhsd, statsmodels v0.11.1) were performed to determine which digestion time points were significantly different (see Fig. 2D).

#### Analysis of Expanded and Unexpanded Mitochondria

To determine if the proExM protocol affects our ability to resolve and quantify mitochondria, we analyzed the number and size of expanded mitochondria compared to unexpanded mitochondria. Tissue from an adult male EGFP reporter mouse was immunostained with COX4 to label mitochondria and expanded using our optimized ExM protocol with overnight digestion in proteinase K (see Expansion Microscopy methods). An adult male tdTomato reporter mouse was similarly processed and immunostained with COX4 but not expanded. Confocal images were taken of both samples for image analysis in FIJI. Note that the COX4 signal was imaged in different color channels for the expanded and unexpanded samples (546nm vs 488nm, respectively).

Confocal images were imported into Fiji and converted to 8-bit for the analysis. An intensity threshold was chosen separately for each image which best represented the signal in the original image. Using the reporter label as a guide, four cells were analyzed from the expanded sample and five cells from the unexpanded sample. Cells were chosen using the same criteria as for the digestion time course analysis. An ROI for each cell was drawn by fitting an oval to the cell body using the oval selection tool, rotating if necessary, and the signal outside of the cell’s ROI was removed for the analysis. The “*nucleus counter”* plug-in from the FIJI “Cookbook” microscopy analysis collection was used to segment mitochondria within the ROI of each cell analyzed. The size threshold used for the expanded sample was adjusted for expansion factor, which was calculated using the ratio of the cell body diameters in the expanded images compared to the unexpanded images. The calculated expansion factor for the expanded images was 3x in the X and Y dimensions-- an expansion factor of 9x in total area.

Once the mitochondria were properly segmented, FIJI’s *“measure “* feature was used to measure the number, area and intensity of each individual mitochondrial ROI. A custom Python code was written to calculate and plot the averages of the mitochondria number, size and total area for the expanded cells and the unexpanded cells. To account for the effect of expansion on size, the area of each mitochondria was divided by the area of the soma to get a normalized mitochondria area. The mitochondrial coverage was calculated by dividing the total summed mitochondrial area in the cell by the area of the cytoplasm *(soma area – nucleus area)* and converting it to a percentage.

### Image acquisition

Images were acquired on an upright Zeiss 710 or inverted Zeiss 700 confocal microscope equipped with a motorized stage, 488/561/633 laser lines, and 5X/0.16 NA, 10X/0.3NA, 20X/1.0 NA water immersion, 20X/0.4 NA air, or 63X/1.4NA lenses. Gels were expanded by washing with 0.001X PBS three times and by incubating overnight at RT in 0.001X PBS. Gels were immobilized in 50mm glass bottom wells (WillCo Wells, HBSB-5040) by applying 2% agarose to the edges of the gels. Gels were then imaged in a fully expanded state in or 0.001X PBS. A subset of images (Fig. 4K-L, Fig. 5G-H) were acquired on an Olympus SpinSR10 spinning disk confocal with a 25X silicone lens. Scale bars are not adjusted for expansion factor unless specifically stated.

### ExM image processing

Czi files were imported into Fiji, individual channel images were split and adjusted for brightness and contrast equivalently across conditions, then images were viewed with the volume viewer plugin (v. 2.01.2). If images showed considerable shift or too many neurons overlapped, a subset stack was created without the offending Z sections. Mode was set to max-projection and interpolation was set to tricubic smooth with z-aspect and sampling optimized per image (typically 0.5-2.0 for each parameter). Snapshots were taken in the XY plane at 1.0 and 2.0 scale in grayscale and 1D. The resulting snapshot images were imported into Photoshop (v. 21.2) and converted to 300 dpi. Any further brightness and contrast edits done in photoshop were minimal and applied equivalently to all comparable images in the figure.

### Semi-quantitative histological assessment of ExM protocols

To quantify the brightness and the quality of the staining after expansion with the various ExM protocols tested in Figures 4 and 5, we developed a scoring system similar to what was done by the Boyden lab (Yu et al, 2020, see Fig. 11). The mean fluorescence of the cropped image was measured in Fiji for each condition, for either the COX4 channel (Fig. 4) or the GOLGA5 channel (Fig. 5) and the reporter channel (GFP or RFP). We normalized the fluorescence by the brightest image in the set to get a fluorescence scale from 0-1. In addition to fluorescence, we gave each image a quality score from 0-1, which took into account how much noise there was in the image and also how close the observed staining was to the expected staining pattern in non-expanded tissue. The fluorescence and quality scores were averaged together and then multiplied by 100 to get an overall score from 0-100 for the antibody and the reporter for each condition.

### RMS analyses

Horizontal 40μm GFP+ sections were immunostained for GFP and GOLGA5 as described above for the pre-IHC-ExM with 8 hours proteinase K digestion. The sections were transferred to a glass bottom plate under a drop of 1XPBS and imaged at 5x (0.16 NA) and 63x (1.4 NA). 63X images were taken at the anatomically identifiable CA1-CA2 border. Sections were then washed in PBS and processed for ExM as described above. After gelation, resulting hydrogels were trimmed and the size of the tissue and gel were measured with a caliper. Tissue and gel sizes were also measured following digestion and after expansion in water. All caliper measurements were taken at the widest point of the section.

Gels were post-stained with DAPI (1:5,000 in water) for 30 minutes and transferred to glass-bottom plates. Gels were then washed in 0.001X PBS overnight. The next day, 10x tile scans of the hippocampus were acquired for each sample. Single 20x images at the CA1-CA2 border were acquired to match the 63x pre-expansion images.

Images were imported into Fiji and processed to obtain matching ROIs from 63X pre-expansion and 20X post-expansion images. Using GFP labeled gels as landmarks, images were cropped to a square which included only overlapping regions in both pre- and post-expansion images (typically 50 x 50 μm in pre-expansion dimensions). Substacks were created from the cropped ROIs to include only the matching cells. Max-projected ROIs were converted to 16-bit grey-scale TIFF files. The pre-ExM image was then scaled in X-Y (with Fiji) to match the pixel dimensions of the post-ExM image, which is needed for proper image registration^38^. The processed images were saved as RAW image files and the associated MHD metadata files were manually created for each image. The GOLGA5 channel was analyzed to get the RMS error. The post-ExM image (moving; Fig. 6H) was registered to the pre-ExM image (fixed; Fig. 6G) using Elastix^39^, with a rigid and then a non-rigid registration as described previously (Chozinski et al, Chen et al). In the first step, a similarity registration was done to align the post-ExM image to the pre-ExM image without warping. The resulting image was then registered again to the pre-ExM image with the B-spline non-rigid registration. The results of the first rigid registration and the second B-spline registration were imported into the Wolfram analysis notebook provided by Chozinski et al. to generate a vector field map of the distortion between the two images (Fig. 6I).

The transformix command was used in Elastix to apply the B-spline transform to a skeletonized image of the post-ExM image. To skeletonize the image, a median and gaussian filter were both applied and the image was binarized. With the output of transformix, the RMS error was then calculated for points along the skeletonized image in Wolfram. For plotting, length was binned per micron and cut off at 30 microns to match previously published RMS error data. The three biological replicates were averaged together (Fig. 6F).

### Statistical analyses

Statistical analyses were done in python (v 3.7) with an alpha of 0.05 considered significant.

## Declarations

### Ethics approval and consent to participate

All animal use and procedures were approved by the Animal Care and Use Committee of Virginia Tech.

### Consent for publication

Not applicable.

### Availability of data and materials

Please contact author for data requests.

### Funding

Research reported in this publication was supported by the National Institute of Mental Health of the NIH under award R00MH109626 to S.F. and by startup funds provided by Virginia Tech. The funders had no role in the design of the study and collection, analysis, and interpretation of data and in writing the manuscript.

### Competing interests

The authors declare that they have no competing interests.

### Authors’ contributions

Conceptualization, S.F.; Methodology, L.A.C., K.E.P., N.A.S., D.L., S.F.; Formal Analysis, L.A.C., K.E.P., N.A.S., S.F.; Investigation, L.A.C., K.E.P., N.A.S., D.L., S.F.; Writing - Original Draft, L.A.C., K.E.P., S.F.; Writing - Review & Editing, L.A.C., K.E.P., S.F.; Visualization, L.A.C., K.E.P., S.F.; Supervision, S.F.; Funding Acquisition, S.F.

## Acknowledgements

We thank the members of the Farris lab, in particular Daniela Gil, for providing feedback and critically reading this manuscript, as well as the Virginia Tech animal care staff for their support.

## Notes

### Competing Interest Statement

The authors have declared no competing interest.

